# Bypassing Striatal Learning Mechanisms Using Delayed Feedback to Circumvent Learning Deficits in TBI

**DOI:** 10.1101/2023.12.27.573465

**Authors:** Ekaterina Dobryakova, Tien T. Tong, Olesya Iosipchuk, Anthony Lequerica, Veronica Schneider, Nancy Chiaravalloti, Joshua Sandry

## Abstract

**Objective:** Feedback facilitates learning by guiding and modifying behaviors through an action-outcome contingency. As the majority of existing studies have focused on immediate presentation of feedback, the impact of delayed feedback on learning is understudied. Prior work demonstrated that learning from immediate and delayed feedback employed distinct brain regions in healthy individuals, and compared to healthy individuals, individuals with traumatic brain injury (TBI) are impaired in learning from immediate feedback. The goal of the current investigation was to assess the effects of delayed vs. immediate feedback on learning in individuals with TBI and examine brain networks associated with delayed and immediate feedback processing.

**Setting:** Non-profit research organization.

**Participants:** Twenty-eight individuals with moderate-to-severe TBI.

**Design:** Participants completed a paired-associate word learning task while undergoing MRI. During the task, feedback was presented either immediately, after a delay, or not at all (control condition).

**Main measures:** Learning performance accuracy; confidence ratings; post-task questionnaire, blood-oxygen-level dependent signal.

**Results:** Behavioral data showed that delayed feedback resulted in better learning performance than immediate feedback and no feedback. In addition, participants reported higher confidence in their performance during delayed feedback trials. During delayed vs. immediate feedback processing, greater activation was observed in the superior parietal and angular gyrus. Activation in these areas has been previously associated with successful retrieval and greater memory confidence.

**Conclusion:** The observed results might be explained by delayed feedback processing circumventing the striatal dopaminergic regions responsible for learning from immediate feedback that are impaired in TBI. Additionally, delayed feedback evokes less of an affective reaction than immediate feedback, which likely benefited memory performance. Indeed, compared to delayed feedback, positive or negative immediate feedback was more likely to be rated as rewarding or punishing, respectively. Findings have significant implications for TBI rehabilitation and suggest that delaying feedback during rehabilitation might recruit brain regions that lead to better functional outcomes.

## Introduction

Learning impairment is a common consequence of moderate-to-severe traumatic brain injury (TBI) that negatively impacts many activities of daily living and leads to overall reductions in quality of life. Because learning forms a critical foundation for successful rehabilitation^1,2^, learning deficits following TBI can exert a negative impact on post-injury outcomes. In the present research we draw on the empirically supported cognitive neuroscience literature to evaluate how TBI impacts learning under different temporal contexts and the associated neural mechanisms.

For a person trying to learn a new task, knowing how successful they are, that is, knowing the outcomes of their actions, constitutes a critical component of the learning process and is a prime example of learning through feedback. An outcome of an action (i.e., feedback), whether positive or negative, allows one to modify their behavior to achieve a specific goal^3^ and helps in acquiring new skills. Feedback varies according to valence (positive or negative) and timing (immediately following the behavior or later [delayed]). Positive feedback indicates to a person that they should continue the action being performed, while negative feedback indicates that they should modify the action to obtain the desired outcome. This type of action-outcome contingency learning or learning through feedback involves the basal ganglia (BG), a deep gray matter structure that can sustain damage in TBI^4^. Along with differences between positive and negative valence, feedback can either be presented immediately following learning or after a delay. Importantly, neuroimaging findings reveal that immediate and delayed feedback processing depend on different BG regions^5^.

Many studies investigating learning through feedback have focused on understanding immediate feedback. When feedback is immediate, there is no delay between the action and the outcome. Research has shown the striatum (part of the BG) to be involved in this type of learning^6–11^. Specifically, human neuroimaging studies in healthy participants have shown that blood-oxygen-level dependent (BOLD) activity in the striatum increases during immediate positive feedback presentation (e.g., positive performance feedback or monetary gain) and decreases during immediate negative feedback presentation (e.g., negative performance feedback or monetary loss)^9^. In this way, the striatum seems to differentially activate depending on the type of feedback. The manner in which a TBI may impact the neural mechanisms associated with learning through feedback has only recently begun to be investigated^12^.

There have been no investigations into the impact of TBI on learning from delayed feedback and similarly, no investigations into the corresponding neural circuitry. However, research in Parkinson’s disease (PD) has demonstrated impaired learning through immediate feedback but preservation of delayed feedback learning^13–15^. The benefits of learning from delayed feedback have been shown with a delay interval as short as 6s from response to feedback presentation in PD^13^, as well as with 25-minute delay interval in healthy individuals^5^. Similar to the deficit in learning from immediate feedback reported in PD^13^, we recently showed that individuals with TBI have a specific deficit in learning through feedback when it is presented immediately after task performance^12^. Learning from immediate feedback is thought to be largely dependent on striatal dopamine function. Numerous animal studies showed that during learning, striatal dopamine neurons exhibit phasic responses to the presentation of a rewarding outcome (feedback)^6^. This dopamine-dependent reinforcement learning is sensitive to the duration of the interval between stimulus presentation and reward (feedback) delivery. If this duration is short (1-2s), dopamine neurons respond as expected to outcome presentation. However, after longer delays, even in the order of seconds, dopamine neurons’ response to rewarding feedback can be as large as their response to unpredicted reward, indicating that dopamine neurons are no longer able to learn the association between the action and the outcome^16,17^.

While immediate feedback seems to largely depend on the striatum, learning from delayed feedback engages a different neural mechanism that does not involve the striatum^5,13,18^. Specifically, there is evidence that learning from delayed feedback recruits the hippocampus in PD^13^ or the more posterior part of the BG, the lentiform nucleus (putamen and globus pallidus) in healthy controls^5^. Given that persons with TBI show a deficit in benefiting from immediate feedback^12^, the provision of delayed feedback might be more effective.

If the cognitive and neural mechanisms associated with delayed feedback are preserved or less impacted in TBI, providing delayed feedback may serve as an efficient means to improve learning following TBI with implications for maximizing progress in rehabilitation. In the current study, we aimed to test whether presenting feedback after a delay would lead to better learning in individuals with TBI. We additionally sought to identify brain regions relied upon by individuals with TBI during learning through delayed feedback.

To address these goals, we presented participants with TBI with a paired-associate word learning task used in Dobryakova and Tricomi’s work^5^. That previous investigation showed that healthy individuals learn equally well from immediate and delayed feedback and, importantly, that these two types of learning involve activation of distinct brain regions. Thus, in the current study, we aimed to evaluate 1) whether learning from delayed feedback leads to improved learning compared to learning from immediate feedback and 2) whether delayed feedback learning results in activation of the lentiform nucleus^5^ and related brain regions in individuals with TBI.

## Methods

### Participants

Thirty individuals diagnosed with moderate-to-severe closed-head TBI were consented to participate in this study. Due to computer malfunction, data from 28 participants were useable for analysis. Consistent with criteria set by the federally funded TBI Model System Program^19,20^, moderate-to-severe TBI was defined as a mechanical blow to the head resulting in at least one of the following: posttraumatic amnesia [PTA] >24 hours, trauma-related intracranial neuroimaging abnormalities, loss of consciousness exceeding 30 minutes, or the Glasgow Coma Scale score of <13 in the emergency department. The information to define injury severity was acquired through reviewing medical records. All individuals were at least one-year post-injury representing the chronic phase. Participants were excluded if they self-reported a significant psychiatric/neurological history (hospitalization or treatment), were taking dopaminergic mediations, or if they were left-handed. See Table 1 for demographic information. The IRB approved this study, and all participants were compensated $150.00 for their participation.

**Table 1.** Demographic information.

| Variable | M (SD) |
| --- | --- |
| Sex |  |
| Male (n) | 18 |
| Female (n) | 10 |
| Race/Ethnicity |  |
| Caucasian (n) | 15 |
| Hispanic (n) | 6 |
| African American (n) | 2 |
| Asian (n) | 2 |
| Multiracial (n) | 3 |
| Age | 38.61 (12.98) |
| Education | 14.36 (2.83) |
| TBI Severity |  |
| Moderate (n) | 12 |
| Severe (n) | 16 |
| Years Since Diagnosis | 7.36 (4.99) |
| M: mean; SD: standard deviation |  |

### Procedure

#### Neuroimaging session

Participants underwent a brain magnetic resonance imaging (MRI) scan conducted on a Siemens Skyra 3T scanner. A T1-weighted pulse sequence was used to collect structural images in 43 contiguous slices (1 × 1 × 1 mm voxels) tilted 30° from the AC-PC line ^21^. Forty-three functional slices were collected using a single-shot echo EPI sequence with 172 acquisitions per run (TR = 2,500 ms, TE = 25 ms, FOV = 192 mm, flip angle = 80°). E-Prime software was used to present the stimulus and for behavioral data collection.

#### Behavioral paradigm

We utilized a paired-associate word learning paradigm, wherein participants were asked to learn unrelated word pairs. This paradigm has been adapted from the prior feedback study conducted by Dobryakova and Tricomi^5^ which consisted of a Study Phase, two Feedback Phases, and a Test Phase (Figure 1). On each trial of the Study Phase, which occurred outside of the MRI scanner, participants were presented with three words on a computer screen: a single target word was presented at the top with two-word options underneath. One of the options was highlighted in green, indicating to the participant that this option is the correct match for the target word. Participants were instructed to memorize the target word and the associated, highlighted option. The words used in the experiment contained 4–8 letters and 1–2 syllables, had Kucera-Francis frequencies of 20–650 words per million, and had high imageability ratings (score of over 400 according to the Medical Research Council database^22^). The words on each trial were matched for word length and frequency. Words presented within the same trial were not semantically related, with a score of less than 0.2 on the Latent Semantic Analysis similarity matrix^23^, and did not rhyme or begin with the same letter. Because processing speed deficits are common in TBI^24^, each trial was presented for 4 sec to allow time for mnemonic association of the word pairs. Trials were presented in random.

**Figure 1.**
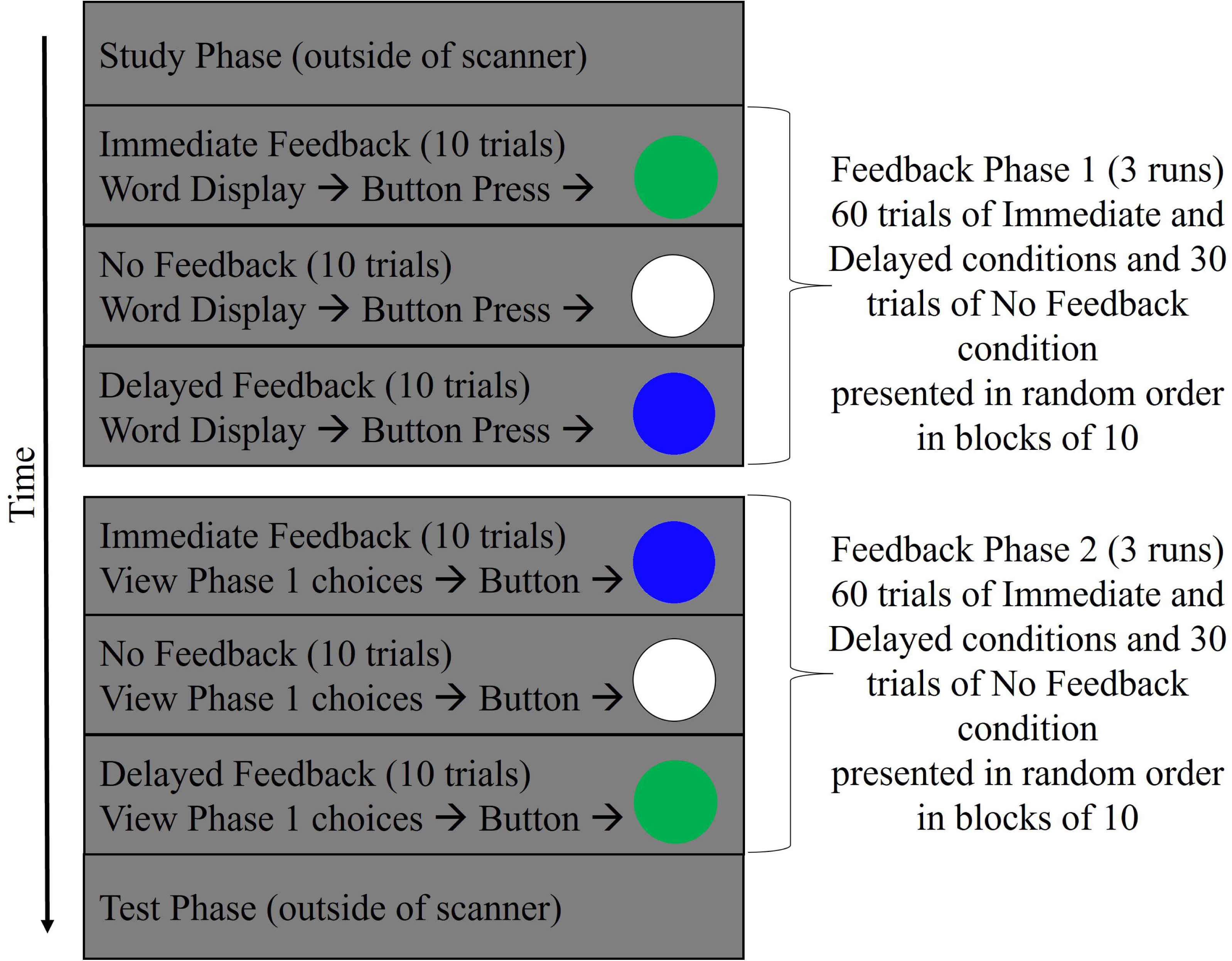
Experimental design of the paired-associate word learning task.

The words from the Study Phase were randomly assigned to the Immediate Feedback (60 trials), Delayed Feedback (60 trials), and No Feedback (30 trials) conditions and were presented in the MRI scanner during Feedback Phase 1 and Feedback Phase 2 (Figure 2). Overall, in Feedback Phase 1, participants completed the first memory test for all conditions, but they only received feedback on their performance for trials in the Immediate Feedback condition. In Feedback Phase 2, participants were asked to respond by pressing their index finger to view their answers in Phase 1. During Feedback Phase 2, participants only received feedback on their performance for trials in the Delayed Feedback condition.

**Figure 2.**
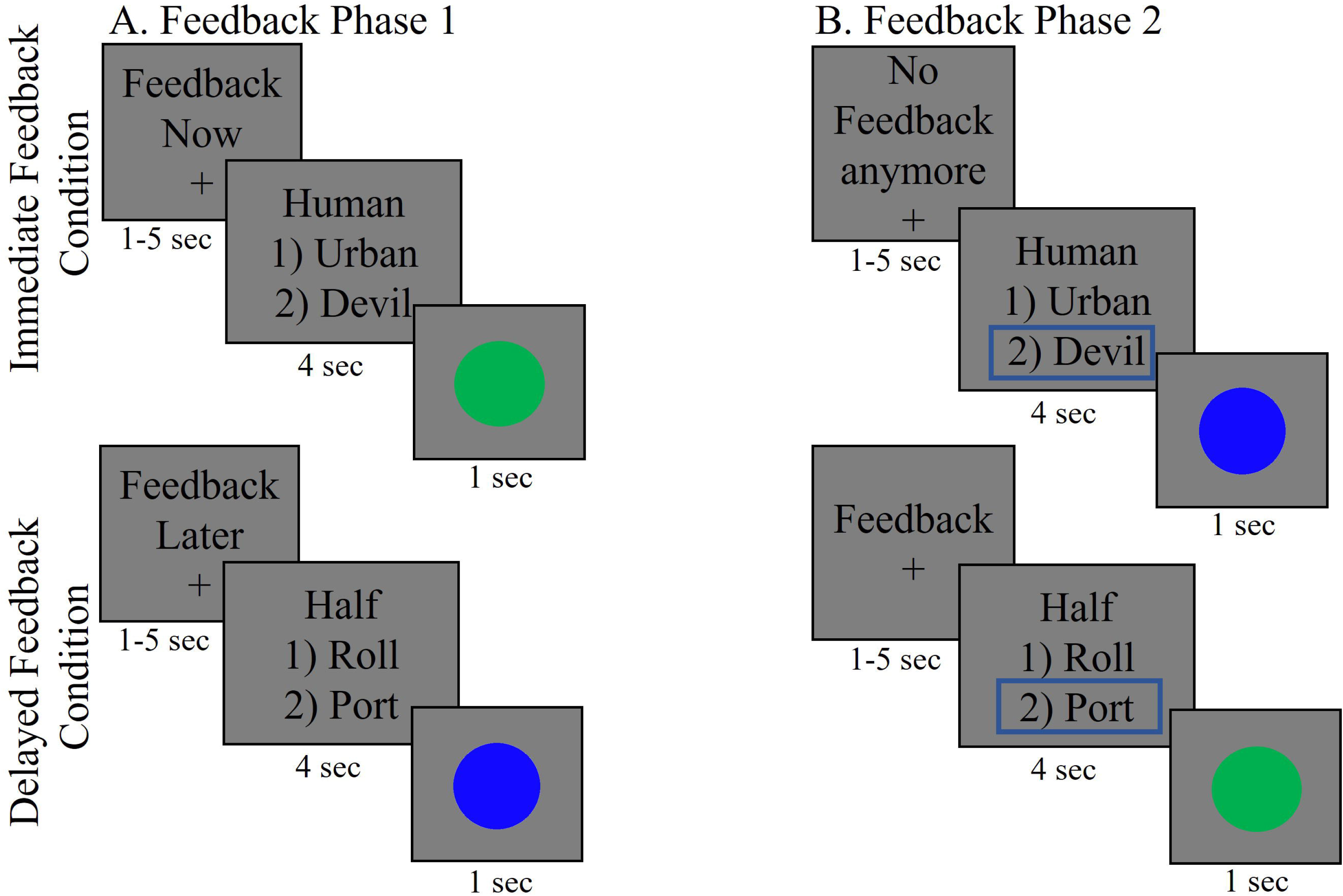
Depiction of a trial in Feedback Phases. Feedback Phase 1: A target word is presented with 2 options underneath; participants select the option they remember to be the correct match; performance feedback is shown for the immediate feedback condition (green circle: correct; red circle: incorrect); delayed and no feedback conditions do not have meaningful feedback (blue circle: delayed feedback; white circle: no feedback). Feedback Phase 2: A target word is presented with 2 options underneath; participants press index finger to view their responses from Phase 1 (highlighted in blue); performance feedback is shown for delayed feedback condition (green circle: correct; red circle: incorrect); immediate and no feedback conditions do not have meaningful feedback (blue circle: immediate feedback; white circle: no feedback)

Specifically, during Feedback Phase 1 (Figure 2A), participants saw a target word along with two options and were asked to select the option that matched the correct paired associate from the Study Phase. Following each response, participants were presented with a feedback screen (1 sec). For the Immediate Feedback condition, the feedback screen showed either a green circle, if participants correctly remembered the word match, or a red circle, if participants incorrectly remembered the word match. For the Delayed and No Feedback conditions there was no performance feedback, instead, participants were presented with a blue and white circle for the respective conditions. Trials were separated from each other with a jittered fixation point (1-5 secs). The conditions were presented in randomized order in blocks of 10 trials.

After Feedback Phase 1, participants began Feedback Phase 2 (Figure 2B). Now, on each trial, participants were presented with trials from Feedback Phase 1, together with a reminder of the option that they previously selected highlighted in blue (for all conditions). To control for the motor response, participants had to press a button unrelated to any word option before being presented with a feedback screen (1 sec). For stimuli in the Delayed Feedback condition, the feedback screen showed either a green circle if participants correctly remembered the word match, or a red circle if participants incorrectly remembered the word match during Feedback Phase 1. The resulting delay between the action (performed during Feedback Phase 1) and the feedback (presented during Feedback Phase 2) was approximately 25 minutes. For trials in the Immediate and the No Feedback conditions, participants were again presented with a blue and white circle, respectively. Trials were separated from each other with a jittered fixation point (1-5 secs). As in Feedback Phase 1, the conditions were presented randomly in blocks of 10 trials.

During the final Test Phase that occurred outside of the scanner, the studied paired associate words were presented in random order, and participants were asked to indicate the correct answer. Each trial lasted 4 seconds and was followed by a confidence-rating question, where participants were given an unlimited time to indicate how certain they were about their response on the scale from 1 to 7 (1=complete guess; 7= completely sure).

At the end of the experiment participants completed a post-task questionnaire where they were asked to indicate how engaging each type of feedback was on a scale from 1 (bored/sleepy) to 7 (very engaging) and which type of positive and negative feedback was more rewarding and punishing, respectively (see Table 2 for exact wording).

**Table 2.**

| Questions on a 7 point scale | I was very<br>bored or<br>sleepy (1) | 2 | 3 | In between<br>(4) | 5 | 6 | I was<br>very<br>engaged<br>(7) |
| --- | --- | --- | --- | --- | --- | --- | --- |
| How engaging did you find the Immediate Feedback portion (first half) of the MRI? | 1 | 0 | 0 | 2 | 4 | 9 | 7 |
| How engaging did you find the Delayed Feedback portion (second half) of the MRI? | 2 | 3 | 6 | 5 | 4 | 1 | 2 |
| *data for 5 people is missing |  |  |  |  |  |  |  |

| Dichotomous Questions | Immediate Feedback<br>(count) | Delayed Feedback<br>(count) |
| --- | --- | --- |
| What type of learning is more difficult? | 14% (4) | 86% (24) |
| Which type of negative feedback felt more punishing? | 54% (15) | 46% (13) |
| Which type of positive feedback felt more rewarding? | 86% (24) | 14% (4) |

## Data analysis

### Behavioral data

To evaluate gains from each type of feedback, we calculated a benefit score by subtracting Feedback Phase 1 accuracy from Test Phase accuracy for each condition and applied a paired-sample t-test. Confidence ratings were analyzed with paired-sample t-test.

### fMRI data

FMRI data processing was carried out using FEAT (FMRI Expert Analysis Tool) Version 6.00, part of FSL (www.fmrib.ox.ac.uk/fsl). Registration of the functional data to the high-resolution structural data was performed using boundary-based registration method^25^. High resolution structural data to MNI2mm standard space registration was performed using FLIRT and further refined with FNIRT nonlinear registration^26,27^. Visual inspection prior to data processing confirmed no significant structural abnormalities that would interference with spatial normalization. The following pre-statistics processing were applied: motion correction using MCFLIRT^28^, non-brain removal using BET^29^, spatial smoothing using a Gaussian kernel of FWHM 6.0mm, grand-mean intensity normalization of the entire 4D dataset by a single multiplicative factor, and high-pass temporal filtering (Gaussian-weighted least-squares straight line fitting, with sigma=45.0s). Time-series statistical analysis was carried out using FILM with local autocorrelation correction^30^. The time series model included regressors corresponding to the 1sec time period of feedback presentation convolved with the double-gamma hemodynamic response function, and their temporal derivatives. A separate regressor was used for each high motion TR, determined using fsl_motion_outliers according to FD > 0.09^31^. Extended motion parameters were included as regressors of no interest (24 regressors: 6 motion parameters, derivatives of the original motion parameters, and the squares of the original and derivatives). One participant did not respond during Feedback Phase 2 and was thus excluded in all group tests that involved Phase 2 contrast.

The second level analysis, which averaged contrast estimates over runs within each subject, was carried out using a fixed effects model by forcing the random effects variance to zero in FLAME (FMRIB’s Local Analysis of Mixed Effects)^32–34^. Group analysis was carried out using FLAME stage 1. To correct for multiple comparisons, Z statistical images were thresholded using a cluster threshold of z>2.3 and a corrected cluster significance threshold of p<0.05^35^.

To address the main hypothesis of the study and examine activation associated with learning through immediate and delayed feedback in individuals with TBI, we ran the following contrast: [(delayed feedback Phase 2 – immediate feedback Phase 2) – (delayed feedback Phase 1 – immediate feedback Phase 1)]. We also explore differences between positive and negative feedback presentation for each feedback type. Thus, the following contrasts were performed: Immediate positive feedback>Immediate negative feedback, Delayed positive feedback>Delayed negative feedback.

## Results

### Behavioral Results

#### Accuracy

Accuracy data were used to calculate the benefit score by subtracting Feedback Phase 1 accuracy from Test Phase accuracy for each condition (see Supplemental Digital Content Figure 1 for percent accuracy of memory performance during Feedback and Test Phases across all conditions). Paired sample t-tests indicated that relative to the No Feedback condition, there was significant benefit of the Delayed Feedback condition, t(27)=3.21, p=.003. In addition, participants performed significantly better on the Delayed Feedback vs. Immediate Feedback condition (t(27)=2.11, p=.04; Figure 3A). No significant differences were observed between the Immediate Feedback and No Feedback conditions (p=.49).

**Figure 3.**
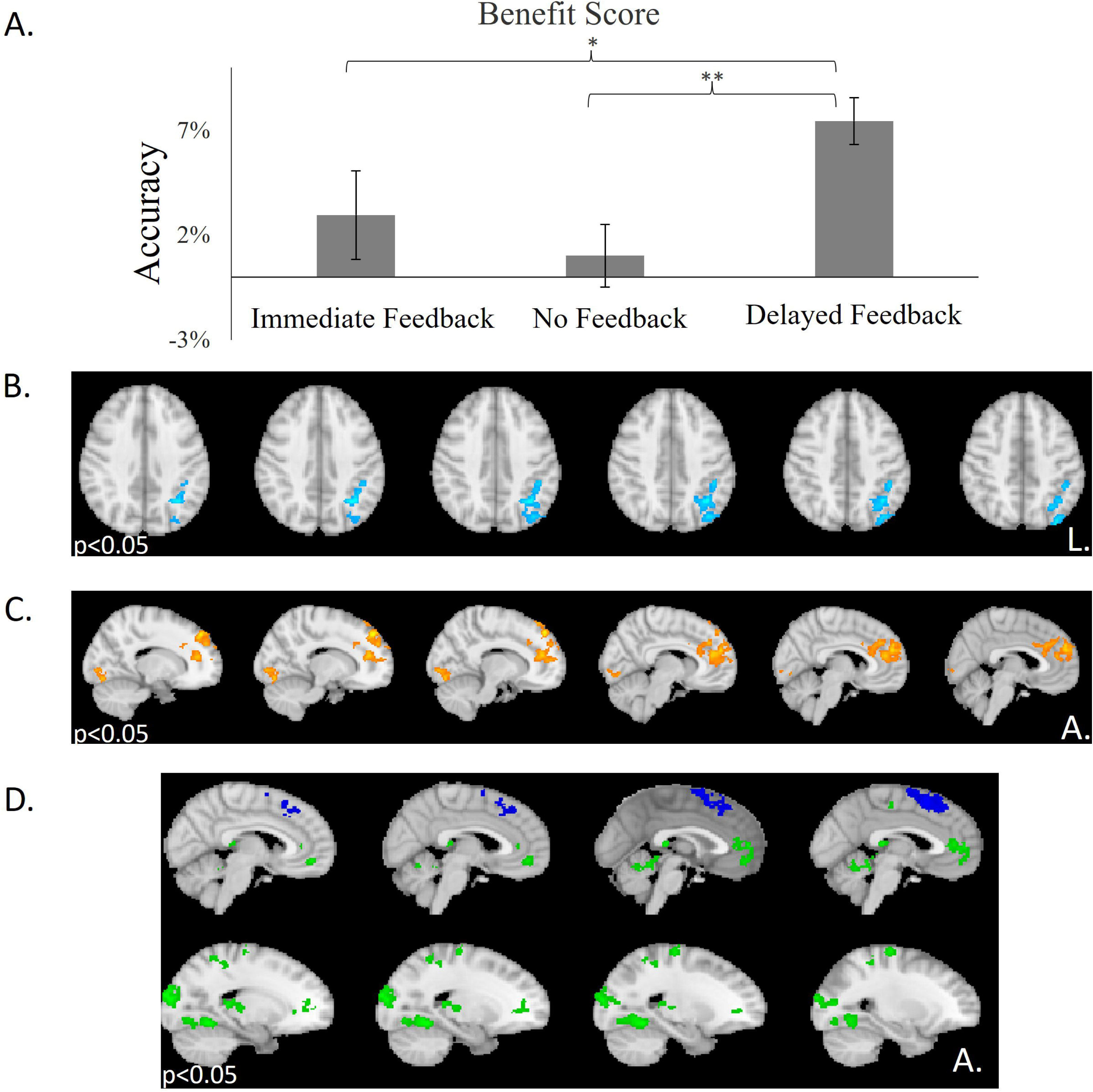
A) Bar graph representing benefit from the 3 Feedback conditions, calculated as the difference between accuracy at Test Phase and Feedback Phase [TestPhase_ACCURACY_ – Feedback Phase_ACCURACY_]. ** indicates p-values less than 0.005; * indicates p-values less than 0.05. B) Delayed feedback presentation was compared with immediate feedback presentation, revealing greater activation in the superior parietal lobe. C) Immediate vs. delayed feedback presentation resulted in activation of the right superior frontal gyrus and left lingual gyrus. D) Brain activation associated with delayed positive vs. delayed negative feedback presentation (green) and delayed negative vs. delayed positive feedback presentation (blue).

#### Confidence Ratings

Two-tailed paired sample t-tests revealed significant differences in confidence ratings with participants being significantly more confident in their responses to words in Immediate Feedback vs. No Feedback condition (t(27)=3.87, p=.001) and in Delayed Feedback vs. No Feedback conditions (t(27)=4.51, p<.0001). Participants were significantly more confident in the responses to words in the Delayed Feedback vs. Immediate Feedback conditions (t(27)=2.54, p<.02).

#### Post-task questionnaires

On the post-task questionnaire, two-tailed Fisher’s exact test showed that significantly more participants indicated immediate positive feedback as more rewarding than delayed positive feedback (p=0.0001). In addition, significantly more participants indicated that learning from immediate feedback was more engaging than learning from delayed feedback (t(22)=6.94, p<0.0001). Furthermore, there was a marginal effect such that those who rated Delayed Negative Feedback as more punishing, performed better on Test Phase Delayed Feedback relative to Immediate Feedback (See additional analyses in Supplementary Digital Content).

### fMRI results

#### Effect Of Delay

To investigate the brain responses to delayed vs. immediate feedback, a whole-brain GLM analysis was conducted to directly compare immediate-and delayed-feedback presentation, while controlling for scanning phase. That is, collapsing across valence, delayed feedback presentation was compared with immediate feedback presentation, while including the presentation trials of the corresponding phase as a control: *[(delayed feedback phase 2 – immediate feedback Phase 2) – (delayed feedback phase 1 – immediate feedback phase 1)*]. This interaction contrast revealed activation in the left superior parietal gyrus. The reverse contrast interaction of immediate vs. delayed feedback presentation [*(immediate feedback phase 1 – delayed feedback phase 1) – (delayed feedback phase 2 -immediate feedback phase 2)]* resulted in activation of the right dorsal ACC extending to dorsomedial superior frontal gyrus and left lingual gyrus (Figure 3B and 3C; Supplemental Digital Content Table 1).

#### Effect of Valence

To gain a better understanding of the effects of valence, we investigated differences in brain activation between positive vs. negative feedback presentation and between negative vs. positive feedback presentation for delayed and immediate feedback conditions. Delayed positive vs. negative feedback presentation resulted in differential activation of the ventromedial prefrontal cortex (VMPFC) and supplementary motor area as well as lingual gyrus (Supplemental Digital Content Table 2). Delayed negative vs. positive feedback presentation revealed activation in the medial superior frontal gyrus and anterior insula (Figure 3D).

For the immediate feedback condition, the contrast between positive feedback presentation vs. negative feedback presentation revealed a large cluster of activation with the peak in the left nucleus accumbens (−14, 8, -12), replicating previous studies^36^. No activation was detected for the opposite contrast of immediate negative vs. positive feedback presentation.

## Discussion

The present study is the first to investigate the influence of the timing of performance feedback on learning in individuals with moderate-to-severe TBI. Here, we show that individuals with TBI learn better from feedback presented after a delay than they do from feedback presented immediately. In addition, we show that, just as in healthy individuals^5^, delayed and immediate feedback processing correspond with different brain regions. These findings highlight how the context of feedback presentation can influence the learning process^9,37,38^ in TBI. This is particularly exciting because it may suggest that incorporating delayed feedback into rehabilitation therapies will lead to long-lasting positive patient outcomes. Cognitive rehabilitation programs that include learning interventions with a feedback component can incorporate delay interval between performance and feedback presentation and this may boost learning. For example, Schmidt et al.^39,40^ asked participants with TBI to prepare a meal during three types of feedback conditions: verbal feedback provided by an OT after meal preparation, video feedback where the video was reviewed with an OT after meal preparation and no feedback. These two investigations showed that participants who were exposed to video feedback had improved self-awareness. The video feedback in this example can be likened to feedback after a delay where individuals were able to review what they did and received feedback about their performance during meal preparation.

One of the differences worth noting compared to the previous findings with healthy individuals^5^ is that contrary to expectations, we did not observe activation of the lentiform nucleus in association with delayed feedback presentation. This might be due to plasticity and reorganization of learning processes due to TBI in the current sample. Such plasticity has been documented in other clinical populations, such as in individuals with PD. Individuals with PD also show impaired learning from immediate feedback^14,15^, however they rely on different brain pathways to process and learn from delayed feedback^13^, specifically the hippocampus.

Similar to individuals with PD, our TBI participants learned better from delayed feedback. Such discrepancy in learning between immediate and delayed feedback and the benefit of delayed feedback might be due to impairment of dopamine transmission during the chronic stages of TBI. Immediate feedback processing depends on dopaminergic structures while delayed feedback processing does not. Evidence from behavioral, pharmacological, and structural MRI studies also point in the direction of impaired dopamine transmission in TBI^41–47^. Animal studies further show that rats given controlled cortical impact injury, a model of closed-head TBI that results in diffuse damage, show altered dopamine transmission^47^ and progressive loss of dopamine-producing neurons. We previously showed that individuals with TBI show deficits in learning from immediate feedback and striatal activation compared to healthy individuals^12^. These studies from various fields of research converge to demonstrate impairment of the dopamine-dependent pathway in TBI and suggest the need for investigating mechanisms and strategies that allow to circumvent this pathway. It is unclear as to why the dopamine system is negatively impacted after TBI but there is ample evidence from previous studies showing that TBI has a great impact on the dopaminergic pathway. The present findings seem to suggest that presenting feedback after a delay circumvents the striatal dopaminergic pathway via evoking greater activation in the superior parietal lobe in individuals with TBI. Delayed feedback in TBI thus seems to rely on compensatory brain mechanisms. Future studies should investigate the degree to which injury severity or other severity-related factors impact the functioning of these compensatory mechanisms and elicit learning through delayed feedback along with comparisons against non-TBI groups.

Another potential explanation for better learning after delayed feedback is that delayed feedback might be causing less of an affective reaction than immediate feedback. Indeed, participants showed higher subjective affective ratings to immediate relative to delayed feedback: positive immediate feedback was more likely to be rated as rewarding. The functional brain activation results also provide converging support for this account. Specifically, the contrast of immediate vs. delayed feedback presentation resulted in activation of the dorsal anterior cingulate cortex and dorsomedial frontal gyrus, prefrontal cortex regions implicated in affective processing^48,49^. Additionally, immediate positive feedback presentation resulted in a large cluster of activation with the peak in the nucleus accumbens, which included VMPFC and rostral anterior cingulate cortex, replicating previous studies of brain activity associated with immediate feedback processing^50,51^. More robust activation in these reward and affective regions was associated with worse performance in the immediate feedback compared to delayed feedback condition. Such hyperactivation of affective regions was also observed in a study where feedback was non-contingent on participants’ actions^52^. The present pattern of activation in combination with prior investigations further support a neural dissociation between immediate and delayed feedback processing. It is possible that the affective nature of immediate feedback presentation “floods” the brain leading to overstimulation that becomes detrimental to learning, at least compared to delayed feedback.

Compared to immediate feedback presentation, delayed feedback presentation resulted in greater activation of regions of the posterior parietal cortex (PPC), specifically, the superior parietal and angular gyrus. Previous research showed that PPC activation during learning is more likely to lead to successful retrieval of information and are associated with greater memory confidence^53^. Indeed, our participants reported higher confidence in their responses to words that belonged to the delayed feedback condition. Further, these PPC regions have also been implicated in active maintenance of information and greater memory capacity^54^. Indeed, the delayed feedback condition requires integration of a previously performed action with feedback, working memory, information and attention organization, as well as retrieval from long-term memory. Thus, activation of these regions may potentially explain why in the current study we observed better learning performance after delayed feedback presentation than immediate feedback presentation. At the same time, delayed positive feedback presentation vs. negative feedback presentation still resulted in VMPFC activation. The VMPFC cluster observed during delayed positive vs. delayed negative feedback presentation, while smaller, overlaps with the cluster observed during immediate positive vs. immediate negative feedback presentation and suggests that delayed positive feedback may still carry subjective affective value but to a lesser degree than immediate positive feedback. These interpretations necessarily require future experimental work to fully unpack how and to what degree affect interacts with feedback processing in TBI.

## Limitations and future directions

It is paramount that we understand how the neural mechanisms involved in learning through feedback are affected in persons with TBI, as such knowledge has the potential to inform rehabilitation practices, influence outcomes, and improve the quality of life of individuals with TBI. While the present investigation demonstrated better learning from delayed compared to immediate feedback, it is important to acknowledge limitations in the generalizability of the findings. Specifically, the generalizability of the results from this study might be limited by the statistical threshold of the fMRI analysis as well as other design considerations, such as learning as measured by recognition, or specific length of the delay between performance and feedback presentation. Still, delayed feedback presented after a short delay^13^ (6s) and long delay interval (25 minutes in the current study) were both beneficial to learning in populations that showed deficits in learning from immediate feedback. Future studies are needed to test potential behavioral and neurological differences between learning after short vs. long delay interval of feedback presentation. Furthermore, as the current study specifically examined learning as defined by recognition performance, future studies should also investigate whether improved learning also generalizes to recall performance. Overall, the dissociation of immediate vs. delayed feedback contingent learning has important implications for rehabilitation where functional outcomes depend on relearning of lost skills after TBI and might suggest that delaying feedback during rehabilitation can lead to better functional outcomes.

## Conclusion

We show that feedback presentation after a delay is more beneficial for learning than immediate feedback presentation. This finding might be explained by the two types of feedback being processed by different brain networks. Moreover, the affective nature of immediate feedback may overwhelm the available neural resources in individuals with TBI and lead to performance decrements. The deficit that is observed in learning from immediate feedback might be due to the impairment of the dopamine pathway after TBI, thus rendering delayed feedback presentation more beneficial. Future studies should further investigate the influence of the feedback timing on learning as well as other contextual factors that can improve learning. More generally, the present findings demonstrate the value of translational cognitive neuroscience in answering clinically relevant-applied questions in TBI.

## Reference

1. Dams-O’Connor K, Gordon WA. Integrating Interventions after Traumatic Brain Injury: A Synergistic Approach to Neurorehabilitation. Brain Impairment. 2013;14(1):51–62. doi:10.1017/BrImp.2013.9

2. Tsaousides T, Gordon WA. Cognitive rehabilitation following traumatic brain injury: assessment to treatment. Mt Sinai J Med. 2009;76(2):173–181. doi:10.1002/MSJ

3. Schlund MW, Pace G. Relations between traumatic brain injury and the environment: feedback reduces maladaptive behaviour exhibited by three persons with traumatic brain injury. Brain Inj. 1999;13(11):889–897. http://www.ncbi.nlm.nih.gov/pubmed/10579660. Accessed April 8, 2015.

4. Huang YN, Yang LY, Greig NH, Wang YC, Lai CC, Wang JY. Neuroprotective effects of pifithrin-α against traumatic brain injury in the striatum through suppression of neuroinflammation, oxidative stress, autophagy, and apoptosis. Sci Rep. 2018;8(1). doi:10.1038/S41598-018-19654-X

5. Dobryakova E, Tricomi E. Basal ganglia engagement during feedback processing after a substantial delay. Cogn Affect Behav Neurosci. 2013;13(4):725–736. doi:10.3758/s13415-013-0182-6

6. Schultz W, Apicella P, Ljungberg T. Responses of monkey dopamine neurons to reward and conditioned-stimuli during successive steps of learning a delayed-response task. Journal of Neuroscience. 1993;13(3):900–913.

7. Apicella P. Leading tonically active neurons of the striatum from reward detection to context recognition. Trends Neurosci. 2007;30(6):299–306. doi:10.1016/j.tins.2007.03.011

8. Apicella P, Scarnati E, Ljungberg T, Schultz W. Neuronal-activity in monkey striatum related to the expectation of predictable environmental events. J Neurophysiol. 1992;68(3):945–960. <GO to ISI>://WOS:A1992JP92500024.

9. Tricomi E, Fiez J a. Feedback signals in the caudate reflect goal achievement on a declarative memory task. Neuroimage. 2008;41(3):1154–1167. doi:10.1016/j.neuroimage.2008.02.066

10. Delgado M, Nystrom LE, Fissell C, Noll DC, Fiez JA. Tracking the hemodynamic responses to reward and punishment in the striatum. Journal of Neuroph. 2000;84:3072–3077.

11. Elliott R, Friston KJ, Dolan RJ. Dissociable neural responses in human reward systems. Journal of Neuroscience. 2000;20(16):6159–6165. <GO to ISI>://WOS:000088676400032.

12. Dobryakova E, Zuckerman S, Sandry J. Neural correlates of extrinsic and intrinsic outcome processing during learning in individuals with TBI: a pilot investigation. Brain Imaging Behav. 2021.

13. Foerde K, Shohamy D. Feedback timing modulates brain systems for learning in humans. J Neurosci. 2011;31(37):13157–13167. doi:10.1523/JNEUROSCI.2701-11.2011

14. Grahn JA, Parkinson JA, Owen AM. The cognitive functions of the caudate nucleus. Prog Neurobiol. 2008;86(3):141–155. doi:10.1016/j.pneurobio.2008.09.004

15. Shohamy D, Myers CE, Grossman S, Sage J, Gluck M a, Poldrack R a. Cortico-striatal contributions to feedback-based learning: converging data from neuroimaging and neuropsychology. Brain. 2004;127(Pt 4):851–859. doi:10.1093/brain/awh100

16. Fiorillo CD, Newsome WT, Schultz W. The temporal precision of reward prediction in dopamine neurons. Nat Neurosci. 2008;11(8):966–973.

17. Kobayashi S, Schultz W. Influence of reward delays on responses of dopamine neurons. Journal of Neuroscience. 2008;28(31):7837–7846.

18. Voermans NC, Petersson KM, Daudey L, et al. Interaction between the human hippocampus and the caudate nucleus during route recognition. Neuron. 2004;43(3):427–435. doi:10.1016/j.neuron.2004.07.009

19. Dijkers MP, Harrison-Felix C, Marwitz JH. The traumatic brain injury model systems: history and contributions to clinical service and research. J Head Trauma Rehabil. 2010;25(2):81–91. doi:10.1097/HTR.0B013E3181CD3528

20. Dijkers MP, Marwitz JH, Harrison-Felix C. Thirty Years of National Institute on Disability, Independent Living, and Rehabilitation Research Traumatic Brain Injury Model Systems Center Research-An Update. J Head Trauma Rehabil. 2018;33(6):363–374. doi:10.1097/HTR.0000000000000454

21. Deichmann R, Gottfried JA, Hutton C, Turner R. Optimized EPI for fMRI studies of the orbitofrontal cortex. Neuroimage. 2003;19(2):430–441. doi:10.1016/s1053-8119(03)00073-9

22. Coltheart M. The MRC psycholinguistic database. Quarterly Journal of Experimental Psychology. 1981;A 33:497–505.

23. Landauer TK, Foltz PW, Laham D. An introduction to latent semantic analysis. Discourse Process. 1998;25(2-3):259–284.

24. Madigan NK, DeLuca J, Diamond BJ, Tramontano G, Averill A. Speed of information processing in traumatic brain injury: modality-specific factors. J Head Trauma Rehabil. 2000;15(3):943–956. http://www.ncbi.nlm.nih.gov/pubmed/10785624

25. Greve DN, Fischl B. Accurate and robust brain image alignment using boundary-based registration. Neuroimage. 2009;48(1):63–72. doi:10.1016/j.neuroimage.2009.06.060

26. Jenkinson M, Bannister P, Brady M, Smith S. Improved Optimization for the Robust and Accurate Linear Registration and Motion Correction of Brain Images. Neuroimage. 2002;17(2):825–841. doi:10.1006/nimg.2002.1132

27. Greve DN, Fischl B. Accurate and robust brain image alignment using boundary-based registration. Neuroimage. 2009;48(1):63–72. doi:10.1016/j.neuroimage.2009.06.060

28. Jenkinson M, Bannister P, Brady M, Smith S. Improved Optimization for the Robust and Accurate Linear Registration and Motion Correction of Brain Images. Neuroimage. 2002;17(2):825–841. doi:10.1006/nimg.2002.1132

29. Smith SM. Fast robust automated brain extraction. Hum Brain Mapp. 2002;17(3):143–155. doi:10.1002/hbm.10062

30. Woolrich MW, Ripley BD, Brady M, Smith SM. Temporal Autocorrelation in Univariate Linear Modeling of FMRI Data. Neuroimage. 2001;14(6):1370–1386. doi:10.1006/nimg.2001.0931

31. Siegel JS, Power JD, Dubis JW, et al. Statistical improvements in functional magnetic resonance imaging analyses produced by censoring high-motion data points. Hum Brain Mapp. 2014;35(5):1981–1996. doi:10.1002/hbm.22307

32. Smith SM. Fast robust automated brain extraction. Hum Brain Mapp. 2002;17(3):143–155. doi:10.1002/hbm.10062

33. Woolrich MW, Ripley BD, Brady M, Smith SM. Temporal Autocorrelation in Univariate Linear Modeling of FMRI Data. Neuroimage. 2001;14(6):1370–1386. doi:10.1006/nimg.2001.0931

34. Siegel JS, Power JD, Dubis JW, et al. Statistical improvements in functional magnetic resonance imaging analyses produced by censoring high-motion data points. Hum Brain Mapp. 2014;35(5):1981–1996. doi:10.1002/hbm.22307

35. Worsley KJ. Statistical Analysis of Activation Images. In: Jezzard P, PM M, Smith S, eds. Functional MRI: An Introduction to Methods. New York, NY: Oxford University Press; 2001:251–270.

36. Dobryakova E, Jessup RK, Tricomi E. Modulation of ventral striatal activity by cognitive effort. Neuroimage. 2017;147(December 2016):330–338. doi:10.1016/j.neuroimage.2016.12.029

37. Tricomi E. Imaging the role of caudate nucleus in feedback processing. 2006.

38. Tricomi E, Fiez JA. Information content and reward processing in the human striatum during performance of a declarative memory task. Cogn Affect Behav Neurosci. 2012;12(2):361–372. doi:10.3758/s13415-011-0077-3

39. Schmidt J, Fleming J, Ownsworth T, Lannin NA. Maintenance of treatment effects of an occupation-based intervention with video feedback for adults with TBI. NeuroRehabilitation. 2015;36(2):175–186. doi:10.3233/NRE-151205

40. Schmidt J, Fleming J, Ownsworth T, Lannin NA. Video Feedback on Functional Task Performance Improves Self-awareness After Traumatic Brain Injury. Neurorehabil Neural Repair. 2013;27(4):316–324. doi:10.1177/1545968312469838

41. Schlund MW, Pace GM, McGready J. Relations between decision-making deficits and discriminating contingencies following brain injury. Brain Inj. 2001;15(12):1061–1071. doi:10.1080/02699050110086887

42. Larson MJ, Kelly KG, Stigge-Kaufman DA, Schmalfuss IM, Perlstein WM. Reward context sensitivity impairment following severe TBI: an event-related potential investigation. J Int Neuropsychol Soc. 2007;13(4):615–625. doi:10.1017/S1355617707070762

43. Newcombe VFJ, Outtrim JG, Chatfield DA, et al. Parcellating the neuroanatomical basis of impaired decision-making in traumatic brain injury. Brain. 2011;134(Pt 3):759–768. doi:10.1093/brain/awq388

44. Shah S, Yallampalli R, Merkley TL, et al. Diffusion tensor imaging and volumetric analysis of the ventral striatum in adults with traumatic brain injury. Brain Inj. 2012;26(3):201–210. doi:10.3109/02699052.2012.654591

45. Tate DF, Wade BSC, Velez CS, et al. Volumetric and shape analyses of subcortical structures in United States service members with mild traumatic brain injury. J Neurol. 2016;33(2):113–122. doi:10.1007/s00415-016-8236-7

46. Bales JW, Kline AE, Wagner AK, Dixon CE. Targeting Dopamine in Acute Traumatic Brain Injury. Open Drug Discov J. 2010;2:119–128. doi:10.2174/1877381801002010119

47. Wagner AK, Sokoloski JE, Ren D, et al. Controlled cortical impact injury affects dopaminergic transmission in the rat striatum. J Neurochem. 2005;95(2):457–465. doi:10.1111/j.1471-4159.2005.03382.x

48. Bush KA, James GA, Privratsky AA, Fialkowski KP, Kilts CD. Action-value processing underlies the role of the dorsal anterior cingulate cortex in performance monitoring during self-regulation of affect. PLoS One. 2022;17(8):e0273376. doi:10.1371/JOURNAL.PONE.0273376

49. Kensinger EA, Ford JH. Guiding the Emotion in Emotional Memories: The Role of the Dorsomedial Prefrontal Cortex. Curr Dir Psychol Sci. 2021;30(2):111–119. doi:10.1177/0963721421990081/ASSET/IMAGES/LARGE/10.1177_0963721421990081-FIG2.JPEG

50. O’Doherty J. Contributions of the ventromedial prefrontal cortex to goal-directed action selection. In: Schoenbaum G, Gottfried JA, Murray EA, Ramus SJ, eds. Critical Contributions of the Orbitofrontal Cortex to Behavior. Vol 1239. Oxford: Blackwell Science Publ; 2011:118–129. doi:10.1111/j.1749-6632.2011.06290.x

51. Rushworth MFS, Noonan MP, Boorman ED, Walton ME, Behrens TE. Frontal cortex and reward-guided learning and decision-making. Neuron. 2011;70(6):1054-1069. doi:10.1016/j.neuron.2011.05.014

52. Spirou A, Chiaravalloti ND, Dobryakova E. Corticostriatal Hyperactivation to Reward Presentation in Individuals With TBI With High Depressive SymptomatologyL: A Pilot Study. Journal of Head Trauma and Rehabilitation. 2018. doi:10.1097/HTR.0000000000000482

53. Wynn SC, Nyhus E. Brain activity patterns underlying memory confidence. European Journal of Neuroscience. 2022;55(7):1774–1797. doi:10.1111/EJN.15649

54. Li X, O’Sullivan MJ, Mattingley JB. Delay activity during visual working memory: A meta-analysis of 30 fMRI experiments. Neuroimage. 2022;255:119204. doi:10.1016/J.NEUROIMAGE.2022.119204

