## Supplementary Materials for "Bypassing Striatal Learning Mechanisms Using Delayed Feedback to Circumvent Learning Deficits in TBI"

**Supplemental Digital Content**

**Supplemental Behavioral Analyses**

**Supplemental Digital Content Figure 1.** Boxplots depicting percent accuracy of memory performance during Feedback Phase (black) and Test Phase (gray) across all conditions.

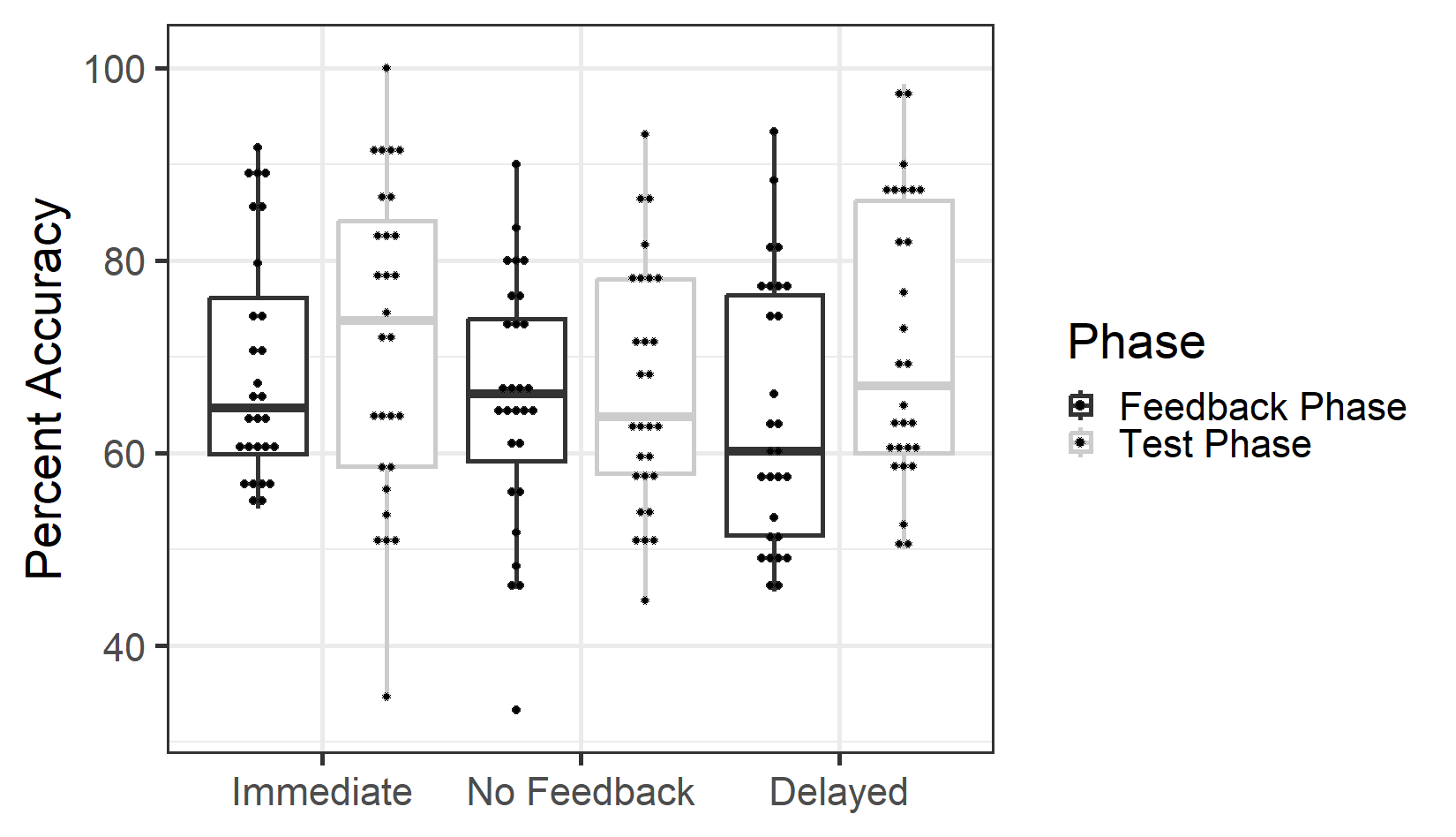

Post-task overall engagement rating was positively correlated with Feedback Phase performance of the No Feedback (*r*(26)=0.40, *p*=.03) and Delayed Feedback (*r*(26)=0.36, *p*=.06, trend) conditions, as well as Test Phase performance of the Immediate Feedback (*r*(26)=0.43, *p*=.02) and Delayed Feedback (*r*(26)=0.32, *p*=.09, trend) conditions. As expected, higher task engagement was associated with better performance across feedback conditions, during both Feedback and Test phases.

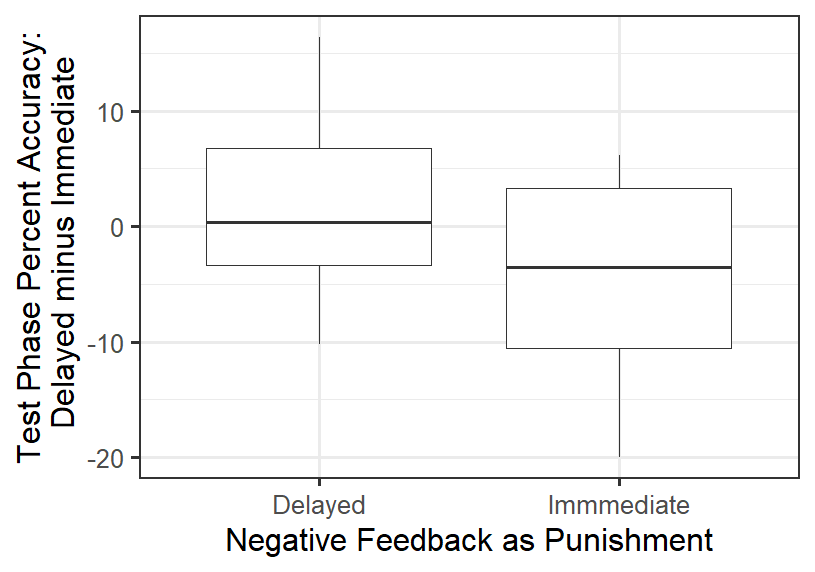

**Supplemental Digital Content Figure 2.** Boxplots depicting Test Phase memory performance (Delayed Feedback minus Immediate Feedback) for people who rated Delayed Negative Feedback as more punishing (left) and those who rated Immediate Negative Feedback as more punishing (right).

Interestingly, there was a marginal effect such that compared to those who rated Immediate Negative Feedback as more punishing, those who rated Delayed Negative Feedback as more punishing performed better on Test Phase Delayed relative to Immediate Feedback (*t*(26)=1.95, *p*=.06, *Supplemental Digital Content Figure 2*). In particular, those who rated Delayed Negative Feedback as more punishing than Immediate Negative Feedback, on average, did not show any differentiation during Test Phase performance between Immediate vs. Delayed Feedback conditions (median=0.33% accuracy difference). On the other hand, there was a trend for better performance in *Test Phase Immediate relative to Delayed Feedback* amongst those who rated Immediate Negative Feedback as more punishing (median=3.19% accuracy difference). In other words, Test Phase performance for the Immediate Feedback condition seemed to be more affective (better performance in a condition in which negative feedback was perceived as punishing) than Delayed Feedback (less differentiation in performance between conditions, even though negative feedback of one condition was perceived as more punishing than the other).

**Supplemental Neuroimaging Results**

SDC Table 1. Effect of Delay

| Delayed vs. Immediate Feedback | |  | |  |  | |  | |  | |
| --- | --- | --- | --- | --- | --- | --- | --- | --- | --- | --- |
| Region | Hemisphere | | Voxels | Peak Z-value | | Peak X | | Peak Y | | Peak Z |
| Superior Parietal Gyrus/Angular Gyrus | L | | 718 | 4.04 | | -30 | | -58 | | 38 |
| Immediate vs Delayed Feedback | |  | |  |  | |  | |  | |
| Region | Hemisphere | | Voxels | Peak Z-value | | Peak X | | Peak Y | | Peak Z |
| Superior and Medial Frontal Gyrus/dorsal Anterior Cingulate Cortex/Frontal Pole/BA9 | R | | 2002 | 4.43 | | 14 | | 46 | | 44 |
| Lingual Gyrus | L | | 724 | 4.04 | | -8 | | -84 | | -10 |

SDC Table 2. Effect of Valence.

| Delayed Positive vs Delayed Negative Feedback Presentation | | |  |  |  |  |
| --- | --- | --- | --- | --- | --- | --- |
| Region | Hemisphere | Voxels | Peak Z-value | Peak X | Peak Y | Peak Z |
| Lingual Gyrus | L | 1375 | 4.27 | -20 | -54 | -12 |
| Left Occipital Pole | L | 853 | 4.55 | -18 | -98 | 18 |
| Precentral Gyrus | L | 796 | 3.66 | -24 | -22 | 68 |
| Ventromedial Prefrontal Cortex (BA 32) | R | 500 | 3.56 | 6 | 48 | -10 |
| Supplementary Motor Area (BA 6) | R | 437 | 3.82 | 54 | 0 | 4 |
| Delayed Negative vs Delayed Positive Feedback Presentation | | |  |  |  |  |
| Region | Hemisphere | Voxels | Peak Z-value | Peak X | Peak Y | Peak Z |
| Superior Frontal Gyrus | R | 2002 | 4.43 | 14 | 46 | 44 |
| Insula | L | 724 | 4.04 | -8 | -84 | -10 |
